# Distinctive BOLD Connectivity Patterns in the Schizophrenic Brain: Machine-learning based comparison between various connectivity measures

**DOI:** 10.1101/084160

**Authors:** Hio-Been Han

**Affiliations:** Department of Psychology, Yonsei University, Seoul, South Korea; Center for Cognitive Science, Yonsei University, Seoul, South Korea; Center for Neuroscience, Korea Institute of Science and Technology, Seoul, South Korea

**Keywords:** schizophrenia, fcMRI, functional connectivity, machine learning

## Abstract

Recent functional magnetic resonance imaging (fMRI) studies have found distinctive functional connectivity in the schizophrenic brain. However, most of the studies focused on the correlation value to define the functional connectivity for BOLD fluctuations between brain regions, which resulted in the limited understanding to the network properties of altered wirings in the schizophrenic brain. Here I characterized the distinctiveness of BOLD connectivity pattern in the schizophrenic brain relative to healthy brain with various similarity measures in the time-frequency domain, while participants are performing the working memory task in the MRI scanner. To assess the distinctiveness of the connectivity pattern, discrimination performances of the pattern classifier machine trained with each similarity measure were compared. Interestingly, the classifier machine trained by time-lagging patterns of low frequency fluctuation (LFF) produced higher classifying sensitivity than the machines trained by other measures. Also, the classifier machine trained by coherence pattern in LFF band also made better performance than the machine trained by correlation-based connectivity pattern. These results indicate that there are unobserved but considerable features in the functional connectivity pattern of schizophrenic brain which traditional emphasis on correlation analysis does not capture.

## Introduction

Functions of the hardly-wired brain require the cooperation of multiple groups of the neuron. There still are some debates about each brain region has its own function (*i. e., modularism*) or only the connections of regions matter (*i. e., connectionism*), there’s no doubt any longer that well-organized brain networks which composed of connections of multiple networks should be operated together to serve complex and integrated brain functions^1^. In line with this idea, recent functional neuroimaging studies in clinical-, systems- and cognitive neuroscience have found abnormalities in anatomical and functional connectivity in the brain with psychiatric disorders who have cognitive deficits, including schizophrenia^2,3^, Alzheimer’s disease^4^, autism spectrum disorder^5,6^, anxiety disorder^7^, obsessive-compulsive disorder^8^, and attention deficit hyperactivity disorder^9^.

Schizophrenia, whose symptoms cannot be easily allocated into single nor few cognitive domains, has been characterized as a model of ‘*disconnected*’ brain^10^. From the level of interneuron wiring^11,12^ to the cortico-cortical circuit wiring^10^, converging evidence now support the idea that pathophysiological mechanisms of altered social, cognitive, and perceptual functions of schizophrenia may be accounted from the abnormalities in local and long-range connections of neuronal circuits at multiple levels. Yet there are still somewhat confounding that distinctiveness of the schizophrenic brain from the healthy brain with the trend of increased or decreased functional connectivity between brain regions, fMRI studies reported altered functional connectivity in the schizophrenic brain, and gently locate this as a putative underlying mechanism of cognitive dysfunction^10,13,14^.

However, since the prominent concept ‘functional connectivity’ in the fMRI studies has been defined as the “*temporal correlation between spatially remote neurophysiological events*” by Friston^15^ and Friston *et al.*^10^, most of the studies dealing with altered functional connectivity in the brain with psychiatric disorder have largely focused on the difference of ‘correlation’ between the regional activities. Indeed, because of the slow nature of blood oxygenation level dependent (BOLD) signal and relatively poor sampling rate of the scanner, less attempts have been made in fMRI studies to describe functional connectivity with other measures of signal similarity with time-frequency domain signal processing between two time-series signals like cross-correlation, coherence and/or synchrony as studies with magneto- or electrophysiological modalities typically have made^16,17^. This raises a question that whether the correlation is able to capture the distinctiveness of network properties in the schizophrenic brain well enough.

Recent decades, there have been attempts to analyze the oscillations from BOLD-fMRI signal with the term ‘Low-Frequency Fluctuation (LFF)’ typically defined in 0.01-0.08 Hz frequency band^18-20^. This resulted in the attempt to define functional connectivity with spectral measures, such as the spectral coherence. In addition, the amplitude in this frequency band of BOLD signal is often related to psychiatric disease such as schizophrenia^21^ and amnestic mild cognitive disorder^22^, describing functional connectivity in the resting-state BOLD signal^23^. However, there’s a lack of evidence about how different the functional connectivity of schizophrenic brain in terms of frequency against the healthy brain.

In this study, the various aspects of distinctiveness in terms of functional connectivity pattern are analyzed in the schizophrenic brain with several time-frequency signal processing techniques. Thanks to the nature of sinusoidal characteristics of LFF activities, the more various aspects of functional connectivity between brain regions can be analyzed. First, functional connectivity can be defined from cross-correlation function. Using this, one can obtain correlation coefficient as a function of the time lag of one signal relative to another. For example, if two signals share exactly same shape of fluctuation but have fixed time lagging with *τ* continuously, cross-correlation function should have its maximum coefficient value (*ρ*) 1 at lag *τ*. So functional connectivity between two time-series signal can be defined as *τ*, the time-lagging pattern when their correlation coefficient gets maximal, and/or *ρ* at lag *τ*, the maximum correlation coefficient value to represent their possible similarity. Second, functional connectivity can also be defined with coherence at specific frequency band. If two time-series BOLD signals share the amplitude of given frequency component continuously, one can assume that two signals share some information which is being represented through cross-spectrum of the frequency channel. With same logic, functional connectivity can be defined as phase synchrony if the signals are somewhat sinusoidal. If two time-series BOLD signals share the phase properties (0 to 2*π*) while oscillating, one can call that two signals are synchronized. In this study, functional connectivity patterns defined by parameters described above (*τ*, *ρ*, at lag *τ*, coherence, phase synchrony) and by the typical correlation value (*ρ* at lag *τ* = 0) are directly compared, to answer the question about distinctiveness of functional networks in the schizophrenic brain.

The aim of this study is to describe the distinctiveness of the functional connectivity in the schizophrenic brain. However, each parameter used here reflects different aspects of signal similarity, it may be too complex to compare the pattern differences directly. To make things simpler and see the distinctiveness of network properties in schizophrenic brain easily, pattern classification based on supervised machine learning technique can be adopted. As machine classifier performs classification totally based on the difference of training dataset, the distinctiveness of schizophrenic brain network can be measured as successful discrimination performance of classifier machine that supposed to detect schizophrenic brain among the unlabeled BOLD connectivity patterns. If one parameter of functional connectivity measure represents the distinctiveness of schizophrenic brain network better than others, discrimination performance of the machine classifier trained by the parameter would be higher than others. By doing that, the distinctiveness of network properties in the schizophrenic brain can be understood better.

## Methods

### Participants & task

The publicly opened datasets were used. This data was obtained from the OpenfMRI database. Its accession number is ds000115. One hundred and two participants performed letter N-back (0-, 1-, or 2-back) working memory task. Data from some participants are excluded because of poor header value of anatomy image (*N* = 4, subject number 033, 068, 086, 102) or missing data points that restrict the time-frequency analysis (*N* = 1, subject number 081), or just simple missing of dataset (*N* = 3, subject number 023, 025, and 048). Remained dataset of 94 participants was used for the analysis. Schizophrenia (*N* = 23) and their siblings (*N* = 32) are classified as schizophrenia group, and healthy control (*N* = 18) and their siblings (*N* = 21) are classified as healthy control group.

### fMRI preprocessing

The preprocessing steps generally followed original publication of the dataset^24^. At first, anatomy image of each participant was reoriented. For the functional images, the basic steps of preprocessing were performed on SPM12 environment including (1) slice timing correction, (2) realignment to compensate for rigid body motion, (3) registration, (4) segmentation, (5) normalization into MNI space (3-by-3-by-3 mm voxel size), then (6) spatial smoothing with three voxels FWHR using a Gaussian kernel. Then bad frames including first five TRs at each run and in which BOLD value remain constant for more than two subsequent TRs were rejected. To improve signal-to-noise ratio, high-pass filtering at 0.009 Hz were performed and nuisance signal including six rigid body motion and global signals were regressed out.

Thirty-four region of interests (ROIs) across four brain networks were defined as Repovš & Barch^24^. BOLD fluctuations of each ROI (sphere with 15mm diameter from the center coordinate) then extracted for the calculation of functional connectivity. As there are 4 types of networks over 34 ROIs, functional connectivity across networks could be defined as follow; *wDMN* (within default-mode network), *wFP* (within frontoparietal network), *wCO* (within cingulo-opercular network), *wCER* (within cerebellar network), and *bDMN-FP* (between DMN-FP networks), *bDMN-CO* (between DMN-CO networks), *bDMN-CER* (between DMN-CER networks), *bFP-CO* (between FP-CO networks), *bFP-CER* (between FP-CER networks), *bCO-CER* (between CO-CER networks).

### Connectivity calculation

All of the functional connectivity calculation were performed between BOLD fluctuations averaged from multiple ROIs on MATLAB R2015a with signal processing toolbox (Mathworks, MA, USA) and custom-written scripts.

#### Correlation & Cross-correlation

Correlation coefficients as a function of the time lag between two ROI BOLD fluctuations were computed from −10 TRs to +10 TRs lag. The coefficient at lag 0 was obtained for typical correlation value, and the coefficient which has the maximum absolute value and the time lag at the point were obtained separately. Coefficients were then transformed into Fisher’s normalized z value. Same procedures for these three parameters were performed to LFF (band-pass filtered with 6^th^ order Butterworth, low-passed then high-passed, at 0.01-0.08 Hz) as well.

#### LFF Coherence

By applying Fast Fourier transform (FFT) with 32 TRs sliding time window, cross-spectral power density in 0.01-0.08 Hz band were obtained then normalized by the signals’ spectral power.

#### Phase Synchrony

To compute the synchronization of ROIs, phase locking value was obtained between ROI BOLD fluctuations which are filtered at 0.01-0.08 Hz band. The phase locking value for the given pair of ROIs was calculated as follow; 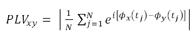, where *ϕ*_*x*_(*t*_*j*_) represents the phase of the sinusoidal BOLD fluctuation of ROI *x* at time *j*^17^.

### Pattern classification

Pattern classification and cross-validation experiments were performed using *Scikit-learn*^25^ the machine learning API on Python 2.7 environment. Non-parametric, supervised learning classifier machine based on decision tree was adopted. The training data were the individual connectivity patterns of all possible combinations between 34 ROIs over 4 types of networks for each task condition, which made the number of input feature dimensions as follow; _34_*C*_2_ (*pairs of ROIs*) × 3 (*task condition*) = 1,623. In another experiment, within- and between-network connectivity patterns were separately used to supervise the classifier (see supplementary materials, Figure S1 and S2).

To validate the classifying performance, the Leave-2-Out procedure was used. As functional connectivity patterns defined with different ways produce different performance, to compare the performance, mean d-prime were obtained as a performance index by averaging the d-primes from 4,371 combinations based on 94 participants’ dataset (_94_*C*_94−2_ pairs) of training-test sets of data in the cross validation procedures. To avoid d-prime going infinite when the probability correct was equal to 0 or 1, maximum (minimum) value of probability correct set to 0.99 (0.01). Scripts for each procedures are available online (https://gist.github.com/Hio-Been, MATLAB script ds115_preproc_v103.m for preprocessing, MATLAB function hb_getConnectivity_ds115.m for obtaining functional connectivity, Python script hb_LPO_ds115.py for machine-learning classification).

## Results

Connectivity patterns defined by correlation of each group and the difference map are shown in Figure 1. As the original article described in Repovš & Barch^24^ found the significant decreased functional connectivity in schizophrenia group for *bFP-CER* and *bCO-CER* network pairs, independent t-test for mean Fisher z values in these pairs produced significant decrease in schizophrenia group, *t*(92) = −2.14, *p* < .01 for *bFP-CER*, *t*(92) = −3.25, *p* < .001 for *bCO-CER*, respectively. Connectivity patterns defined by other parameters are shown in Figure 2. Multiple comparisons of mean parameter values for all possible network pairs go beyond the aim of this study, means of other parameters are not analyzed (Figure S1).

**Figure 1.**
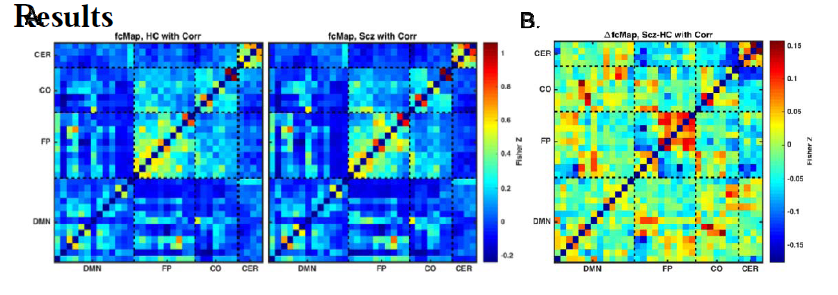
Mean functional connectivity patterns defined by correlation. (A) Result of healthy control group (left) and schizophrenia group (right). (B) Difference between two connectivity patterns. Note that positive value represents higher values in the schizophrenia group, and each network (DMN : Default-mode network, FP : Frontoparietal network, CO : Cingulo-opercular network, CER : Cerebellar network) has different number of ROIs.

**Figure 2.**
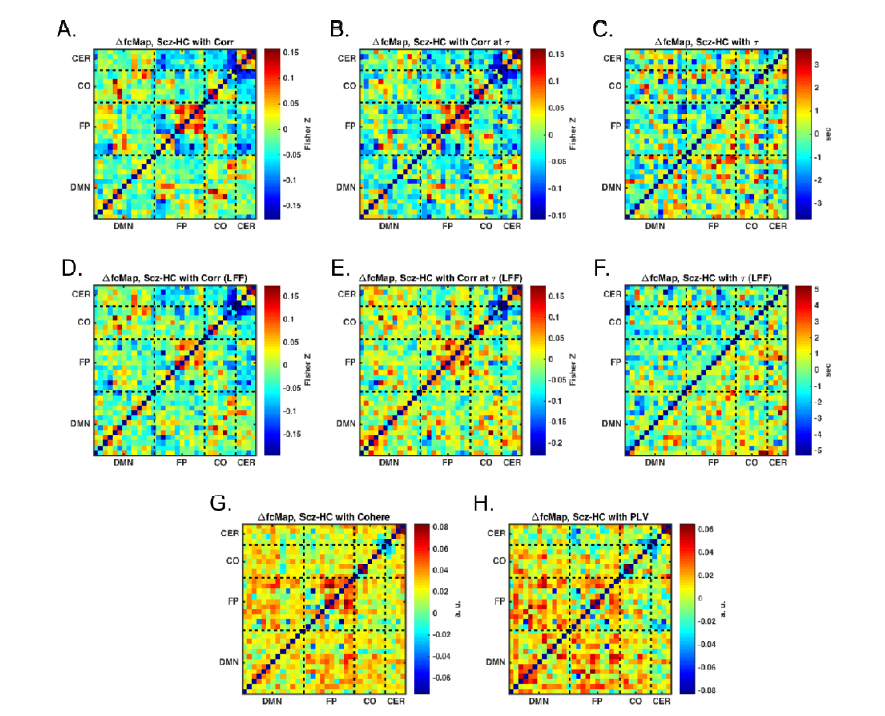
Mean differences in functional connectivity patterns defined by (A) correlation, (B, C) maximum value of absolute correlation coefficient and tau at the point, (D) correlation calculated from LFF band, (E, F) maximum value of absolute correlation coefficient and tau at the point calculated from LFF band, (G) coherence at LFF band, and (H) phase synchrony calculated from LFF band. Note that positive value represents higher values in the schizophrenia group, and each network (DMN : Default-mode network, FP : Fronto-parietal network, CO : Cingulo-opercular network, CER : Cerebellar network) has different number of ROIs.

To test whether there are significant differences in the mean classifying performances, repeated measures analysis of variance was performed. Mean d-primes and standard error of the mean d-primes are as follow (Figure 2.); ‘Correlation (*M* = 0.650, *SE* = 0.046)’, ‘Correlation at *τ* (*M* = 0.671, *SE* = 0.048)’, ‘*τ* (*M* = 0.987, *SE* = 0.049)’, ‘Correlation (LFF) (*M* = 1.475, *SE* = 0.047)’, ‘Correlation at *τ* (LFF) (*M* = 0.029, *SE* = 0.047)’, ‘*τ* (LFF) (*M*= 1.906, *SE* = 0.044)’, ‘Coherence (*M* = 1.402, *SE* = 0.047)’, ‘Phase locking value (*M* = −0.316, *SE* = 0.046)’. Mean d-primes across conditions were significantly different, *F*(7, 4336) = 767.11, *p* < .001. Post-hoc contrast analysis revealed that there were significant differences for every pair of mean d-primes except 2 pairs (‘correlation’ vs ‘correlation at *τ*’, *p* = .32; ‘correlation (LFF)’ vs ‘coherence (LFF)’, *p* = .20) produced significant differences, all the *p*s < .05. In addition, all conditions except ‘correlation at *τ* (LFF)’ produced non-zero classifying performance, all *p*s < .05. Most interestingly, mean d-prime of the condition ‘corr at *τ* (LFF) ’ produced the highest performance. Further classifications which used only the single network out of 16 network pairs (within- and between over 4 types of networks) did not produced higher than this condition (Figure S2).

**Figure 3.**
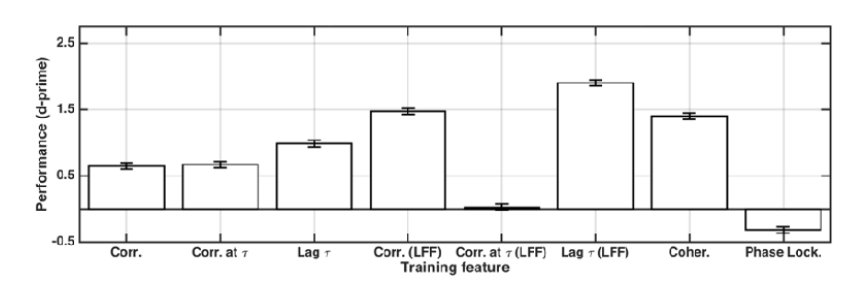
Result of leave-two-out cross validation procedure for the functional connectivity patterns including all within- and between-network combinations. Error bars represent standard errors of the means.

## Discussion

In this study, the distinctiveness of the network property of schizophrenia functional brain networks was compared across several measures of functional connectivity. As a result, it turns out that some parameters defined in time-frequency domain using low-frequency band captured the distinctiveness of schizophrenic brain network better than typical measure defined by correlation. Among them, time-lagging patterns in LFF produced the highest distinctiveness of schizophrenia functional brain networks. At the same time, there was actually no distinctiveness between schizophrenia and healthy brain in the condition of ‘correlation at *τ*’ because the mean d-prime of the machine trained with this parameter was not statistically different from zero, which measures the correlation after compensating the timing differences between two time-domain signals. These indicate that the time lagging should not be compensated to capture the distinctiveness of network property in the schizophrenic brain in the LFF activities.

Considering the nature of coherence, it makes sense there is no difference in the performance of ‘coherence’ and ‘correlation (LFF)’ because they are the same measure conceptually, although there are some practical differences in the filtering method or specific options in transformation parameters. However, the negative value of mean d-prime observed in the ‘phase locking value’ condition is difficult to interpret. As there is no previous report of phase synchronization in this band BOLD oscillations, it will be an important future work to demonstrate the synchronization properties in LFF.

To sum up, all of these results suggest that one should consider time-frequency activities when to examine the functional connectivity patterns in BOLD signal in the schizophrenic brain better. This is the first time, to my best knowledge, to compare the distinctiveness of functional connectivity patterns defined by time-frequency parameters of schizophrenia relative to the healthy brain directly. In addition, there was no previous attempt to define functional connectivity pattern as a function of time-lagging value (*i. e.*, *τ*) *per se.* Since the results of this study show the time-lagging value *τ* is much more robust feature to capture the distinctiveness of altered functional connectivity in the schizophrenic brain, a rigorous study about this finding should be located for next step. Advantages of this approach, defining functional connectivity in BOLD signal with time-frequency signal processing techniques, can be easily found to other research themes such as various type of cognitive dysfunction (*e. g*., ADHD) or specific cognitive functions whenever the functional connectivity matters.

## Acknowledgements

I especially thank Prof. Do-Joon Yi of Yonsei University for his great advice to build up the concept of the study, and Prof. Grega Repovš of the University of Ljubljana for sharing the dataset and for his helpful advice about fMRI preprocessing methods.

## Contributions

H.-B.H. designed the study, analyzed and interpreted the data, and drafted the manuscript.

## Additional Information

Competing financial interests: The author declares no competing financial interests.

---

**Figure S1.**
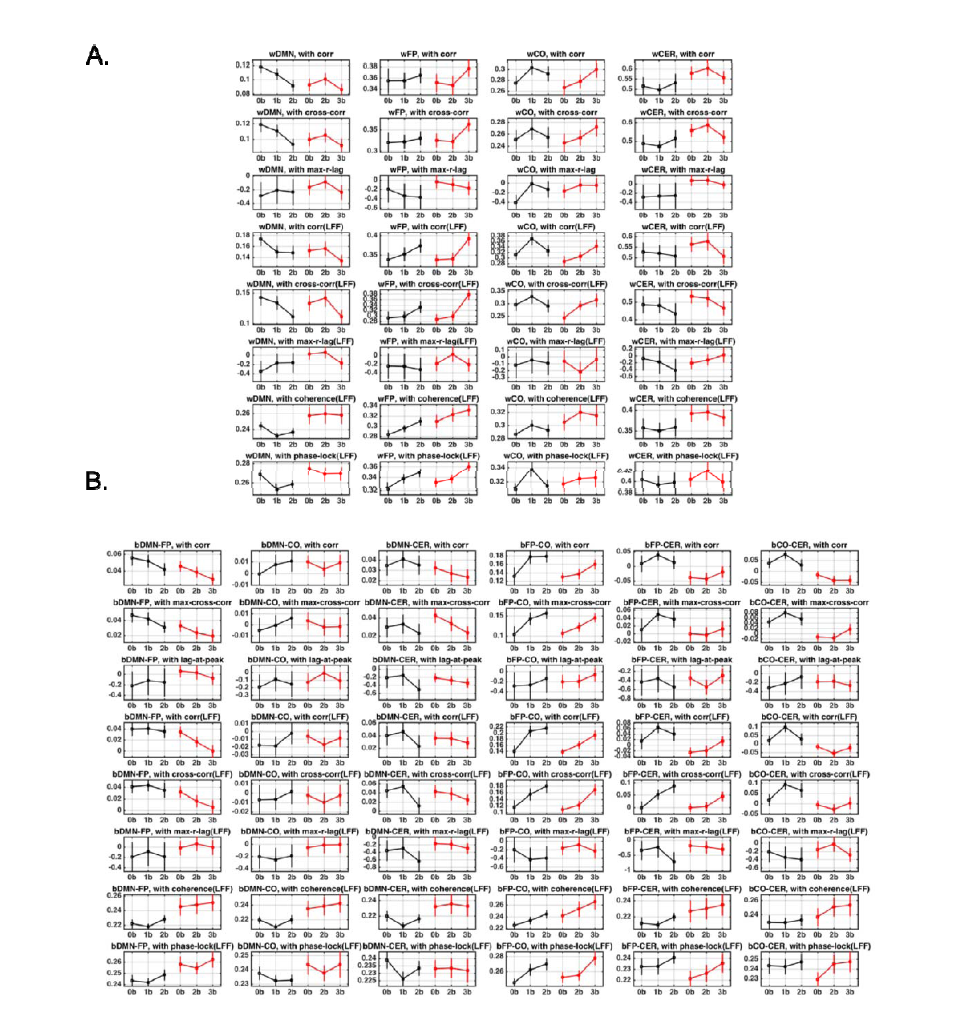
Mean values of connectivity defined by each parameter (A) within network, and (B) between networks (red : schizophrenia, black : healthy control; 0b : 0-back, 1b : 1-back, 2b : 2-back).

**Figure S2.**
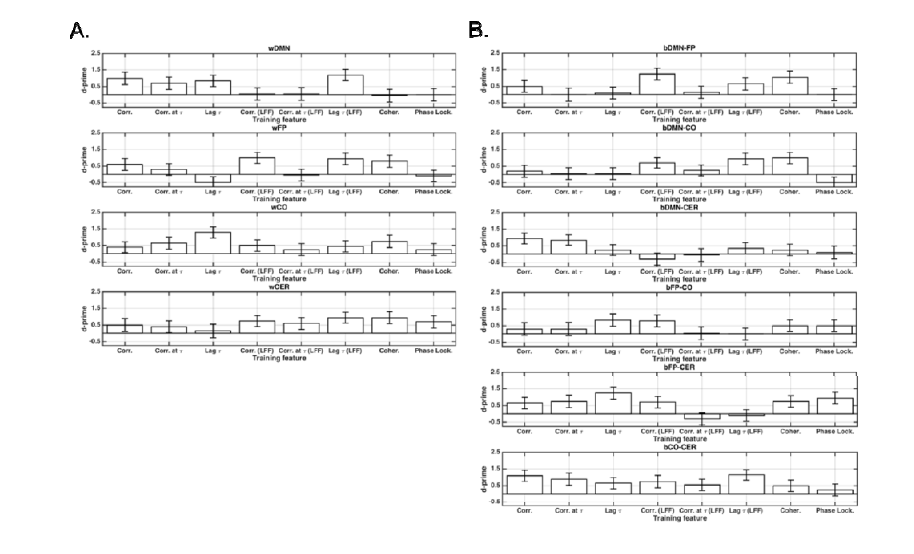
Mean classifying performance defined by single pair of (A) within network, and (B) between networks. Note that error bars represent standard errors of the mean.

